# Who takes care of the kids at when? Sex differences in avian parental care

**DOI:** 10.1101/2021.10.20.465177

**Authors:** Daiping Wang, Wenyuan Zhang, Xiang-Yi Li Richter

## Abstract

Parental care in birds consists of many elaborate forms, including nest building, incubation, and offspring provision, but we still do not know how much each parent contributes to the different forms. Furthermore, the variations, relationships, and potential drivers of sex differentiation in providing care across different care stages remain largely unknown. Here, we surveyed species in birds and uncovered remarkable differences in the sex role patterns across different care forms. This result implied that parental care should not be treated as a unitary trait but as a composite of integrated features with great variations. Further analyses revealed moderate correlations of the sex roles between care forms, indicating the existence of shared intrinsic drivers. We tested the effects of sexual selection, certainty of paternity, predation risk, and offspring’s life history traits in driving sex role variations. Results showed that species with strong sexual selection on males or uncertainty of paternity tend to have female-biased care.

## Introduction

Birds often provide extensive parental care that enhances their offspring’s survival and future reproductive fitness. Avian parental care comprises diverse forms, including nest building, incubation, provisioning the offspring, and defending them against predators^1–3^. Despite the benefits that parental care brings, it costs energy, time, and the opportunity for extra-pair mating and/or starting a new clutch. Furthermore, it may increase the predation risk of the parents. Consequently, there are conflicts between the male and female parent and between parents and offspring. In cooperative breeding species, the conflicts involve helpers of different relatedness with the breeders and the dependent offspring. These intricate relationships have inspired theoretical studies about the optimal parental care strategies of each parental sex and helpers. Early models characterized parental care as an all-or-none choice between deserting and caring^4–7^, while later models generally treated parental investment as a continuous trait. Most theoretical work, however, treats parental care as a unitary trait rather than a composite of several functionally integrated characteristics. A few rare exceptions have considered task specialization between parents, such as feeding the young and defending them from predators^8,9^, but these models do not make predictions on how parents contribute to different tasks over time across a breeding cycle.

Focusing on one or a small set of species, optimal levels of parental efforts have been studied as functions of various factors, including brood quality^10,11^, the certainty of paternity^12–14^, operational sex ratio and sexual selection^15,16^, and sex-specific life history characters such as adult mortality^17^ and the ability to care^18^. Special attention has been paid to which sex should provide care, yet without distinguishing care forms across the whole breeding cycle^19–22^. Do sex-specific parental strategies differ across distinct care forms? In other words, if one sex has participated in nest building, would it also incubate the eggs laid in that nest and/or feed the chicks after they hatch? In birds, the variations, relationships, and potential drivers of sex differentiation in providing care across different care stages remain largely unknown. To figure out the degree to which sex roles in parental care differ between distinct care forms across a breeding cycle, we first survey the participation of males and females in three typical care forms (i.e., nest building, incubation, and offspring provisioning) across broad avian taxa. We then test three main hypotheses regarding the relationships of sex roles across distinct parental care forms. Finally, we aim to uncover possible driving forces of sex variation in parental care across different forms.

The three main hypotheses focus on the consistency of parental care in temporally consecutive forms. In other words, if a male has built the nest, would he continue to incubate the eggs, or leave the task to the female? Similarly, if a female has incubated the eggs, would she continue to feed the chicks after they hatch, or leave them to the male? Increasing evidence for consistent individual behavior across taxa and social contexts suggested that the inflexibility of behavior (i.e., behavioral consistency) may be beneficial^23–27^. For example, being consistently bolder or more active than others may consistently benefit the growth and fecundity of the focal individual under certain conditions^28^. In the case of parental care, it may be favorable for a sex to specialize and consistently provide care across different forms, which we refer to as the “consistent expertise hypothesis”. However, given that there are often parental conflicts over costly caring efforts, flexible behavioral responses might be more adaptive^29^. Indeed, theoretical^30^ and empirical studies found that males and females can communicate and negotiate their parental effort^31–35^, and the negotiation rules can be sex-specific^36^. Following the logic with a broader view, we hypothesize that each parent may negotiate with their partner regarding who provides parental care across different forms and the stages of breeding. For instance, at the stage of nest building, if the male has built the nest solely, the female may agree to provide care alone in the next stage (i.e., incubation), followed by the male joining her in offspring provision thereafter. We refer to this mechanism as the “complementary negotiation hypothesis”, the second hypothesis we test. Furthermore, it is also possible that the parental efforts of males and females are neither consistent nor complementary, but determined independently by their sex-specific opportunity cost of breeding in different forms of care. Indeed, caretaking males of the black coucals were found to have different success rates in siring extra-pair offspring at varying stages of parental care^37^, suggesting that the opportunity cost of parental care can change over time. We refer to this mechanism of parental effort allocation as the “distinct pattern hypothesis”, which is the third hypothesis that we test.

Besides testing the consistency of parental care, we also aim to identify the driving forces of sex role variations across different care forms. In particular, we test the roles of sexual selection, certainty of paternity, nest predation risk, and offspring’s life history traits in driving the variations. The four potential driving forces are chosen because there are clear theoretical predictions of their effects on sex differences in parental care^38^. Strong sexual selection on males is predicted to produce female-biased care^15,16^, except when females prefer to mate with care-providing males, which can lead to the evolution of male-biased care^39^. Sexual selection has also been shown to associate with evolutionary transitions between major patterns of parental care^40,41^. Another important predictor of parental care investment is the certainty of parentage. The difference between male and female parents in expected parentage (e.g., due to female extra-pair mating) is predicted to produce female-biased care, and males should invest more in caring for their genetic offspring^12–14^. In addition, sex differences in parental care can arise if providing care is more costly or less efficient for one sex than the other. For example, high nest predation risk may select for female-biased care when females have more cryptic plumages than males, which is common in passerines^42^. Being drabber, females may be less likely to attract predators to the offspring and themselves when providing care. Furthermore, the life history traits of offspring are of interest to study because they reflect broods’ reproductive value and needs. Because parents’ caring efforts are linked to the trade-off between their current and future reproductive fitness, they are expected to invest more in broods of higher reproductive value^10,11,43–45^.

We collected parental care data from all 1533 avian species in the Birds of the World database^46^, tested the consistency of sex roles across nest building, incubation, and offspring provision, and identified the main driving forces of the sex role variations at different stages of a breeding cycle.

## Results

### Large variation in sex roles across different forms of parental care

We found remarkable diversity regarding which sex provides care across the three different care forms in birds (Figure 1, N = 1533 species). At the stage of nest building, both partners built the nest in 938 (61.19%) species (including 56 species of cooperative breeding); the female built the nest alone in 538 species (35.09%); the male built the nest alone in only 57 species (3.72%). At the stage of incubation, female-only care (768 species, 50.10%) and biparental care (740 species, 48.27%) were about equally common, while male-only care was very rare (25 species, 1.63%). At the stage of offspring provisioning, biparental care was the dominant form (1344 species, 87.67%, including 139 species of cooperative breeding), followed by female-only care (166 species, 10.83%), while male-only care (23 species, 1.50%) continued to be the rarest sex role pattern.

**Figure 1.**
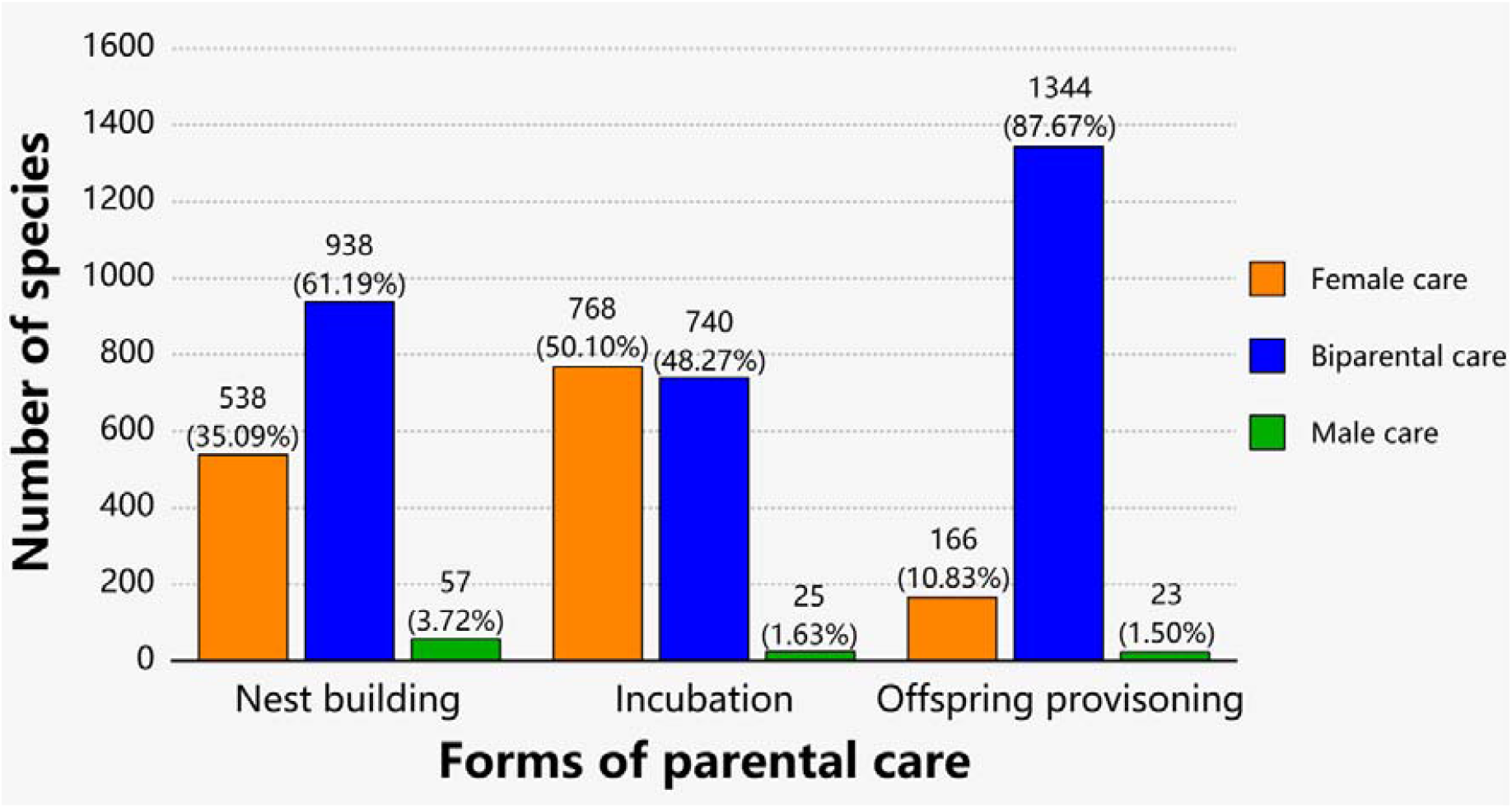
Variation of sex roles in different forms of parental care. The results were based on 1533 species with full data on sex roles (‘Female care’, ‘Biparental care’, and ‘Male care’) across three forms of parental care: nest building, incubation, and offspring provisioning. Bars of different colors represent different sex role categories. Note that a few species of cooperative breeding were grouped into the ‘Biparental care’ category (N = 56 species in nest building, N = 48 in incubation, and N=139 in offspring provisioning).

### Moderate consistency of sex roles across breeding stages

We revealed moderate consistency of sex roles between temporally consecutive stages of parental care. The direct phenotypical correlations of sex roles between nest building and incubation, between incubation and offspring provisioning, and between nest building and offspring provisioning are all positive (Figure 2 and Figure 3). Moreover, the multivariate phylogenetic model showed that the three phylogenetic correlations are also positive (nest building – incubation: 0.509 ± 0.051; incubation – offspring provisioning: 0.419 ± 0.050; nest building – offspring provisioning: 0.348 ± 0.022, Figure 3, Table S1). In short, both phenotypical and phylogenetic correlations suggested that sex roles of parental care across different stages are consistent, supporting the ‘consistent expertise hypothesis’ (Figure 3). Additionally, the multivariate phylogenetic model showed that the sex roles across the three reproductive stages have strong phylogenetic signals (λ = 0.800 ± 0.021; 0.902 ± 0.016; 0.769 ± 0.026, respectively, Figure 3, Table S1).

**Figure 2.**
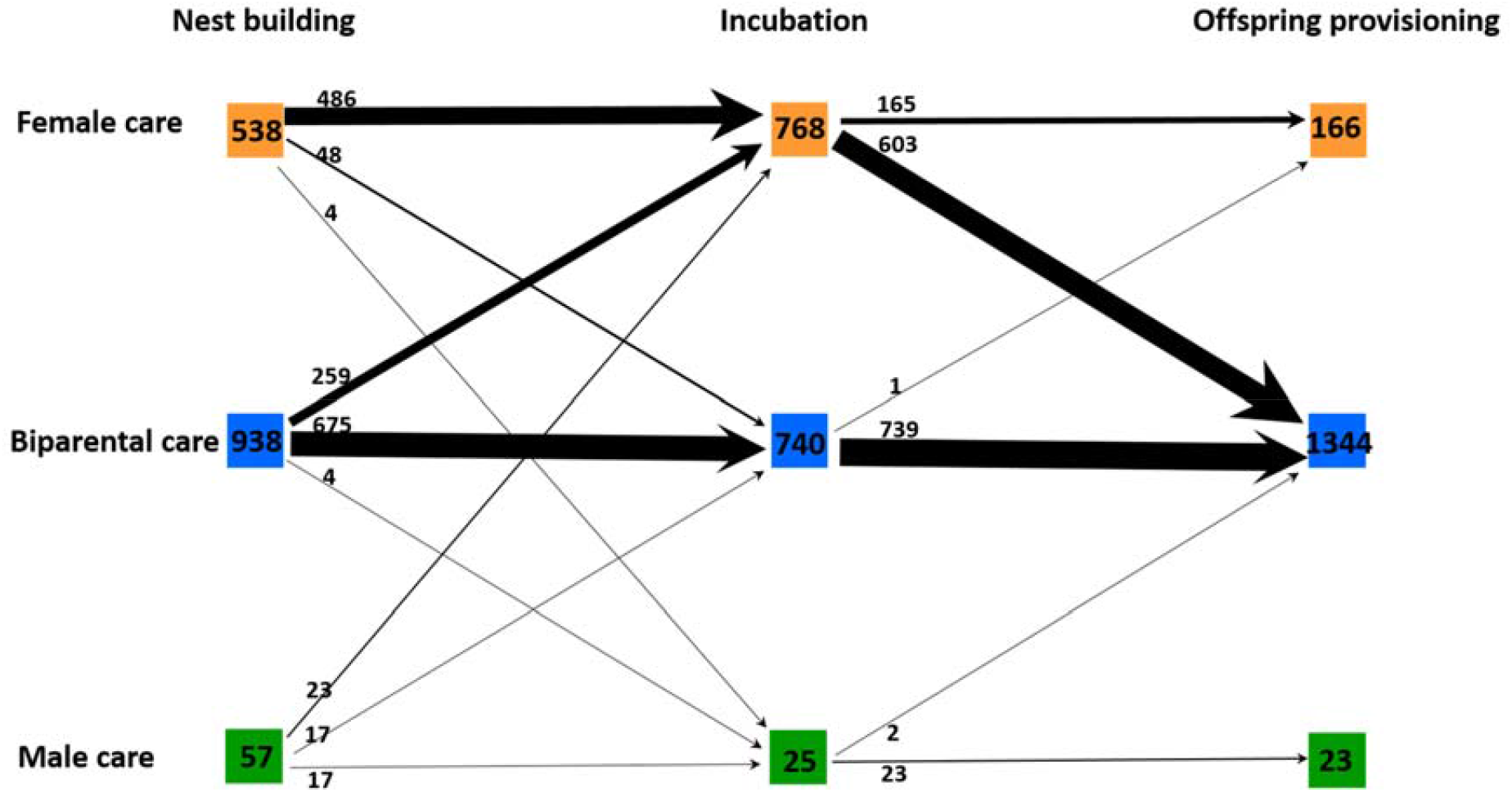
The number of avian species of each sex role pattern (i.e., ‘Female care’, ‘Biparental care’, and ‘Male care’) from nest building to offspring provisioning.

**Figure 3.**
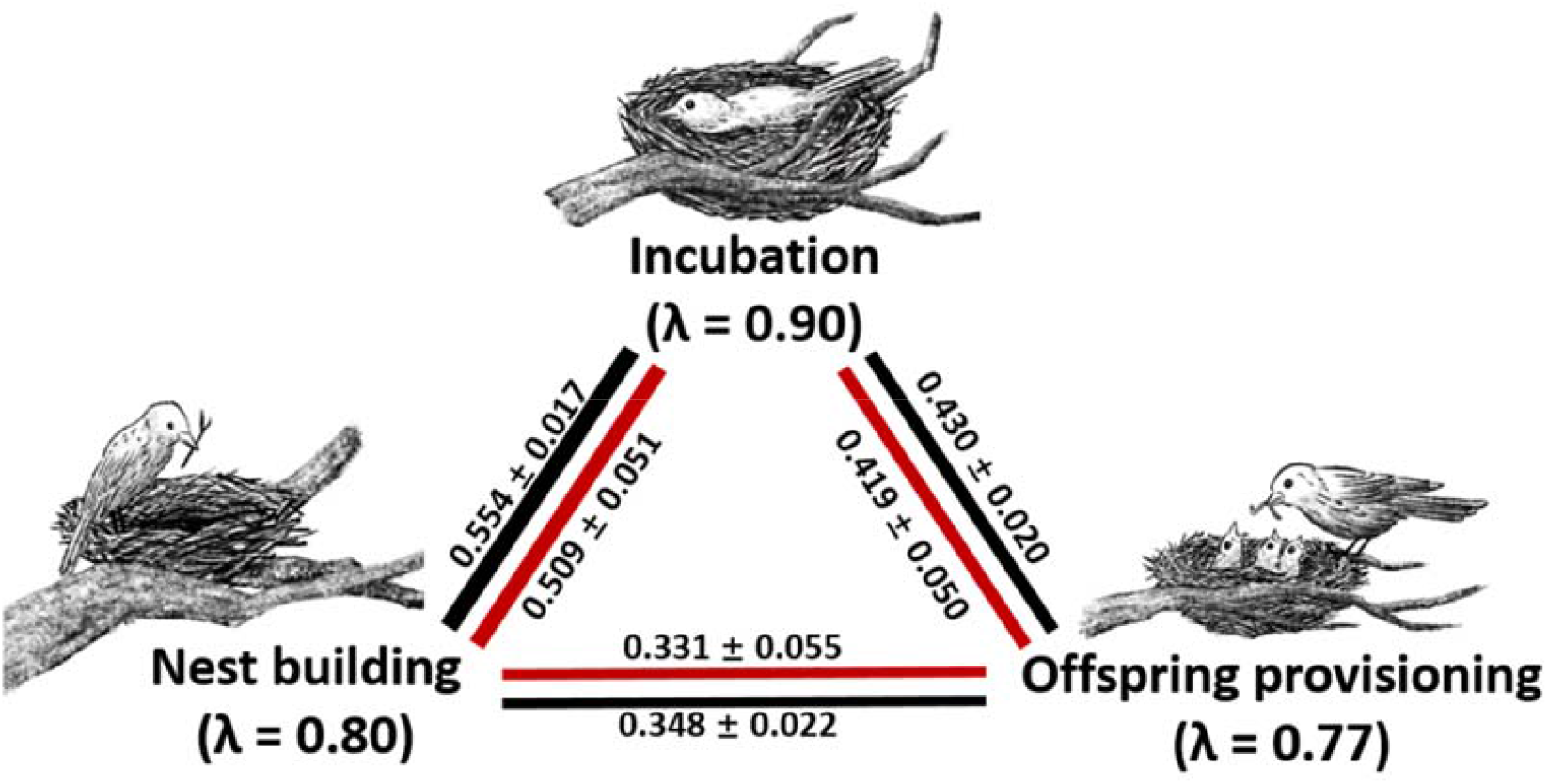
Phenotypical and phylogenetic correlations of sex role patterns between three distinct parental care stages. The phenotypical correlations (black) were estimated based on 1533 species. The phylogenetic correlations (red) were estimated using a multivariate phylogenetic model based on 1050 species.

### Biases towards female care under strong sexual selection

Overall, our statistical analysis of the linear mixed-effect model (Table 1: Model 1) revealed a significant association between sexual selection and the parental roles in nest building, incubation, and offspring provisioning (*t =* −11.5, *p* < 0.001; Figure 4). The consistent pattern of sexual selection is markedly stronger in the ‘Female care’ than the ‘Biparental care’ and ‘Male care’ category across all three forms of care. Furthermore, sex roles in parental care across three different care forms tended to depend on research effort (*t =* 2.2, *p =* 0.03). The random effects of ‘family’ and ‘genus’ explained 31% and 9% of the variation in the response variable, respectively, indicating a moderate phylogenetic signal. The phylogenetic signal was even stronger in the phylogenetically controlled regression model (λ = 0.59, 95 CI: 0.47 – 0.67, Table S2: Model 1). The estimated direction and degree of the fixed effects (i.e., ‘form of care’, and ‘sexual selection’) from the phylogenetically controlled regression model were almost the same as the linear mixed-effects model except for the fixed effect ‘research effort’ which turned to be not significant (compare Table S2: Model 1 with Table 1: Model 1).

**Table 1.** Summary of statistics of five linear mixed-effects models (Model 1 to Model 5). For the random effect, the size of the variance components is shown. The estimate with its standard error (SE), *t*-value, and corresponding *p* value is shown for each fixed effect.

| | | Estimate<br>($\beta \pm SE$) | *t* | *P* |
| --- | --- | --- | --- | --- |
| Model 1 | Random effects: |  |  |  |
|  | Family (n = 131) | 0.065 |  |  |
|  | Genus (n = 484) | 0.019 |  |  |
|  | Residual | 0.126 |  |  |
|  | Fixed effects: |  |  |  |
|  | Intercept | -0.292 ± 0.028 | -10.4 | - |
|  | Nest incubation | -0.176 ± 0.015 | -11.4 | <0.001 |
|  | Offspring provisioning | 0.256 ± 0.015 | 16.5 | <0.001 |
|  | Sexual selection | -0.108 ± 0.009 | -11.5 | <0.001 |
|  | Research effort | 0.00005 ± 0.00002 | 2.2 | 0.03 |
| Model 2 | Random effects: |  |  |  |
|  | Family (n = 64) | 0.01 |  |  |
|  | Genus (n = 124) | 0.104 |  |  |
|  | Residual | 0.141 |  |  |
|  | Fixed effects: |  |  |  |
|  | Intercept | -0.261 ± 0.059 | -4.4 | - |
|  | Nest incubation | -0.175 ± 0.042 | -4.2 | <0.001 |
|  | Offspring provisioning | 0.350 ± 0.041 | 8.4 | <0.001 |
|  | EPP | -0.003 ± 0.002 | -1.7 | 0.09 |
|  | Research effort | -0.00002 ± 0.00005 | -0.4 | 0.72 |
| Model 3 | Random effects: |  |  |  |
|  | Family (n = 65) | 0.113 |  |  |
|  | Residual | 0.146 |  |  |
|  | Fixed effects: |  |  |  |
|  | Intercept | -0.281 ± 0.060 | -4.7 | - |
|  | Nest incubation | -0.185 ± 0.043 | -4.4 | <0.001 |
|  | Offspring provisioning | 0.333 ± 0.042 | 7.9 | <0.001 |
|  | EPBr | -0.0005 ± 0.001 | -0.5 | 0.65 |
|  | Research effort | 0.000 ± 0.000 | -0.2 | 0.86 |
| Model 4 | Random effects: |  |  |  |
|  | Family (n = 55) | 0.034 |  |  |
|  | Genus (n = 138) | 0.011 |  |  |
|  | Residual | 0.121 |  |  |
|  | Fixed effects: |  |  |  |
|  | Intercept | -0.496 ± 0.044 | -11.3 | - |
|  | Nest incubation | -0.241 ± 0.033 | -7.3 | <0.001 |
|  | Offspring provisioning | 0.527 ± 0.033 | 16 | <0.001 |
|  | Nest daily predation rate | -0.149 ± 0.888 | -0.2 | 0.87 |
|  | Research effort | 0.000 ± 0.000 | -0.02 | 0.98 |
| Model 5 | Random effects: |  |  |  |
|  | Family (n = 140) | 0.096 |  |  |
|  | Genus (n = 497) | 0.044 |  |  |
|  | Residual | 0.137 |  |  |
|  | Fixed effects: |  |  |  |
|  | Intercept | 1.562 ± 0.163 | 9.6 | - |
|  | Nest incubation | -0.122 ± 0.017 | -7.2 | <0.001 |
|  | Offspring provisioning | 0.332 ± 0.017 | 19.6 | <0.001 |
|  | Nestling developmental time | 0.129 ± 0.094 | 1.4 | 0.16 |
|  | Clutch size | -0.065 ± 0.038 | -1.7 | 0.09 |
|  | Research effort | -0.00003 ± 0.00003 | -1.3 | 0.19 |

**Figure 4.**
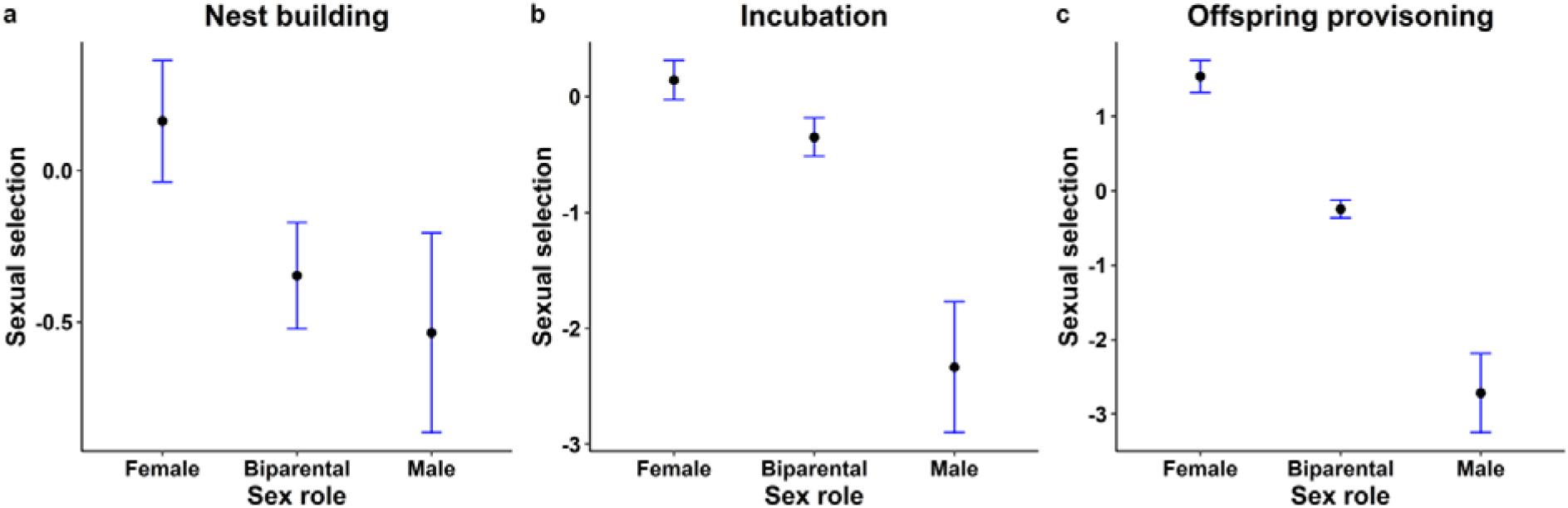
Sexual selection (PC1 of mating system, sexual size dimorphism, and sexual dichromatism) scores of three sex role categories in parental care across three distinct care forms. Plots in each panel showed the mean value with 95% confidence intervals of sexual selection scores in each sex role category.

### Association between the certainty of paternity and male care

Our statistical analyses (Table 1: Model 2 and Model 3) revealed a consistent association between the certainty of paternity and the investment of males in parental care across breeding stages. Species with high levels of extra-pair paternity (EPP) or having broods that contain extra-pair offspring (EPBr) tended to show less male care (i.e., ‘Female care’ and ‘Biparental care’ were more abundant; EPP: *t =* −1.7, *p =* 0.09, EPBr: *t =* −0.5, *p =* 0.65; Table 1: Model 2 and Model 3). In the phylogenetically controlled regression model, the estimated direction and degree of extra-pair paternity (either EPP or EPBr) were stronger and more significant (EPP: *t =* −2.7, *p =* 0.01, EPBr: *t =* −1.5, *p =* 1.5; Table S2: Model 2 and Model 3).

### No clear association between sex roles in parental care and predation risk

Model 4 did not show a significant association between sex roles in parental care and nest daily predation rate across three different care forms (*t =* −0.2, *p =* 0.87; Table 1: Model 4; *t =* 0.6, *p =* 0.53; Table S2: Model 4). Hence, the difference between males and females in the cost of providing care (predation risk in this case) appeared to be non-essential in determining which sex provides care.

### No clear association between the number of carers and offspring’s life history traits

Model 5 showed no significant association between the number of carers in each care form and nestlings’ developmental time (*t =* 1.4, *p =* 0.16; Table 1: Model 5; *t =* 0.6, *p =* 0.56; Table S2: Model 5). There was also no clear association between the number of carers and clutch size (*t =* −1.7, *p =* 0.09; Table 1: Model 5; *t =* −1.3, *p =* 0.18; Table S2: Model 5). These results suggest that the reproductive value of the current brood (represented by clutch size) and brood needs (represented by both clutch size and nestling developmental time) were not determinant factors of the number of individuals that provide care.

## Discussion

Our survey of more than 1500 species of birds revealed that sex roles of parental care differ substantially across different care forms (i.e., nest building, incubation, and offspring provisioning). However, statistical tests showed there is moderate consistency regarding which sex provides care across the three different stages, supporting the ‘consistent expertise hypothesis’. Using five linear mixed-effects models, we further tested several ecological and evolutionary factors that may explain sex differences across different forms of parental care, and we identified sexual selection and the certainty of parental care to be the main driving forces. Uniparental care by females tended to be more frequent in species under strong sexual selection, and males of species with high certainty of paternity were more likely to contribute to parental care. However, we did not find a significant association between nest predation rate and sex-specific contribution to parental care. There was also no evidence that offspring’s life history traits that reflect their reproductive value and brood needs played a role in the number of carers. Our major findings remain unchanged by excluding uncertain species from the dataset (Supplementary Table S3).

### Parental care is not a unitary trait regarding which sex provides care

Our results regarding parental roles at the stage of offspring provisioning are similar to the findings in Cockburn (2006)^47^, which did not include data on the stages of nest building and incubation. The marked differences in sex role patterns across different breeding stages we found in this study (Figure 1) indicate that parental care is not a unitary trait regarding which sex provides care but a composite of several integrated features with great variations. Our results and the lack of theoretical predictions highlighted important knowledge gaps in our understanding of parental care as a package with several functionally integrated traits, and how males and females were selected to fulfill different sex roles in the evolutionary time scale. Studies in birds have identified several factors that affect the (relative) contributions of the male and female parents in the ecological time scale, including the harshness of abiotic environments, especially temperature and rainfall^48–51^, predation risk^52,53^, the vulnerability of offspring in the absence of parental care^54,55^, and the body condition of the parents themselves^56^. Would those factors also play a role in driving sex roles evolution in different care forms in the evolutionary time scale? Do they co-evolve with each other? And how eco-evolutionary feedback may affect the evolutionary trajectories and evolutionary transitions? Future studies in both empirical and theoretical aspects are needed to answer those questions.

### Moderate sex role consistency over parental care

Despite the differences in the overall pattern of sex roles across different forms of parental care, our statistical analysis showed moderate consistency of sex roles between consecutive care forms (Figure 2, Figure 3), supporting the ‘consistent expertise hypothesis’^57^ rather than the two alternatives (i.e., the ‘complementary negotiation hypothesis’ and ‘distinct pattern hypothesis’). This result thus adds to the increasing evidence for consistent individual behaviors across social and ecological contexts^23,26,27^. Although sex role specialization leading to consistency in parental care patterns seems intuitive to expect, the underlying evolutionary mechanisms and whether such specialization provides any consistent fitness advantages are still unclear. Our Model 1 suggests that strong sexual selection on the sex that does not provide care may also contribute to sex role consistency of the opposite sex (Table 1). The parental care patterns in the lekking species and sex role reversed species provide the strongest support for this hypothesis. Males of the lekking species such as grouses, paradises and manakins are under extraordinary sexual selection. They have the most extravagant appearances and courtship displays but contribute nothing but genes to the offspring, leaving females to provide all forms of parental care alone^58,59^. But in sex role reversed species such as jacanas, phalaropes and coucals, where sexual selection on females is stronger than in males, it is the males that are responsible for all forms of parental care duties^60^. Note that the two aforementioned explanations are not mutually exclusive –– sex role specialization of one sex and strong sexual selection on the other sex may synergistically cause the former sex to provide care consistently throughout different stages of parental care^61^.

### Strong sexual selection was tied to sex-biased care of all forms

Our analyses showed a consistent pattern of sexual selection being stronger in species of female-only care (male-only care in sex-role reversed species) than in species of biparental care across three different forms of parental care. This result is in agreement with the Darwin-Bateman paradigm that predicts sexual selection on males leading to the evolution of conventional sex roles^62^, and concurred with a recent survey of 659 bird species from 113 families, which found that parental cooperation decreased with the intensity of sexual selection and skewed adult sex ratios^63^. The study^63^ focused on the association between sexual selection and the “inequality” between males and females in parental care contributions, and therefore they analyzed the parental care data without sex-specificity. Also, despite that the parental care data in the previous study contained eight different parental care activities (corresponding to different care forms in our study), the parental cooperation score was calculated by averaging the statistically centered extent of biparental care across the different activities. In contrast, we surveyed more species (1533 species in total) and associated data on sex-specific contributions of parental care in three distinct forms. The two studies are thus complementary to each other, and the combined results suggest that the role of sexual selection on the evolution of sex-biased parental care may be widespread across avian taxa and across different forms of parental care.

### Uncertainty of paternity selected against male care

Our analysis showed a significant association between extra-pair paternity and reduced male care across different parental care forms, in agreement with previous comparative studies with a smaller number of species^64–67^. Although theories generally predict that males should invest more in the care of their genetic offspring and adjust their parental efforts to their share of paternity in the nest^12–14,61^, empirical supports have been mixed, with abundant exceptions where males do not seem to react to the loss of paternity by reducing their parental care efforts, such as in dunnocks^68,69^, reed buntings^70^, and western bluebirds^71^. Recent theoretical studies revealed some conditions where males may evolve to be insensitive to the loss of paternity, e.g., in cooperative breeding species where offspring help to raise their younger (half-)siblings^72^, or in the presence of male alternative reproductive tactics where the “sneaker” males specialize in gaining extra-pair paternity^73^. Empirical studies also found that in species where males were not sensitive to paternity loss, paternal care may not be costly in terms of parental survival^70^ and/or the loss of opportunities for siring extra-pair offspring^37^. Few comparative studies (for a rare exception, see Griffin (2013)^74^) have tested the role of potential factors that may explain the presence or absence of male response to paternity loss by reducing or withholding paternal care, probably due to a limitation of detailed data on life history traits related to parental care across species. Future efforts in generating and collating such data are therefore indispensable to a better understanding of the relationship between the certainty of paternity and male investment in parental care.

### Nest predation risk did not shape sex roles in parental care

Our analyses did not show a significant association between nest predation risk and sex differences in parental care. This result is surprising as it contrasts with a previous survey of 256 species of passerine birds, which found that the frequency of nest visits decreased as the risk of nest predation increased, as frequent bouts of incubation could increase the visibility of a nest^75^. Similar results were found also in seven species of arctic sandpipers^76^. Given that the plumage of females is usually drabber and more cryptic than males, we expected species with high nest predation to show more female-biased care. The lack of correlation could be due to either anti-predatory adaptations, confounding factors that masked the effect of female cryptic plumage, or a combination of both. Species that endure high nest predation risk may have evolved strategies that minimize activities that could attract predators, like long on- and off-bouts of incubation^48^, and males with brighter plumage may evolve to attend the nest largely at night when visual predators were inactive, such as in the red-capped plover^77^. Confounding factors such as nesting site quality and the shape of nests may also override the advantage of the drabber plumage of females in providing care. For example, a study using 10 species of open-nesting birds in Arizona, USA revealed a positive correlation between nest predation and parental activity only when nest site effects were considered^78^.

### Brood needs and offspring’s reproductive value did not affect the number of care providers

Since broods of larger sizes and longer nestling developmental time generally have higher needs, we expected that more carers (i.e., both parents relative to a single parent, or breeders and helpers relative to only the breeders) were required to provide the elevated amount of care. But no such association was found in our data. Our results suggested that the amount of parental care a brood receives may not necessarily increase with the number of carers. Indeed, models have shown that a parent may or may not compensate for a reduction of parental effort by the other depending on various factors, including the marginal benefit/harm to offspring as a function of total care received, how well each parent is informed about brood needs, and how well the parents can monitor each other’s investment^19,21,22,79^. Negotiation between parents can even produce cases where the offspring do better with one parent than two^20^. Experimental studies by (temporally) removing a parent also showed that the compensation patterns can vary widely from a matching reduction, through no, partial, and full compensation, to even over-compensation^80–83^. Therefore, species are likely to have evolved redundancy in their abilities to provide care, and such abilities could be beneficial to secure reproductive success in cases of losing a partner and/or helper.

## Conclusion

Through a survey of more than 1500 species of birds, we found great diversity in terms of which sex provides care across three different forms (i.e., nest building, incubation, and offspring provisioning). We also found moderate consistency of the sex roles between consecutive stages of care, indicating there might be some shared intrinsic drivers that lead one parent to provide care in different forms. Our models showed that the intensity of sexual selection is the primary driver of the sex-role variations we found in distinct care forms. We also found that uncertainty of paternity selects against male care. As a whole, our results suggest that parental care should not be treated as a unitary trait, but as a composite of integrated features with great variations. Besides those findings, we also identified important knowledge gaps for future theoretical and empirical investigations. For example, we still lack testable theories that make predictions on the relative efforts of male and female parents in different care forms. And we still do not fully understand why males react to a loss of paternity by reducing paternal care in some species but not in others. Would the effects of sexual selection, certainty of paternity, predation risk, and offspring life history traits we found in birds play similar roles in other animal groups? Do other factors, such as adult sex ratio, operational sex ratio, and sex-specific adult mortality, also play a role in shaping sex-role patterns in different forms of parental care? And how do these driving factors interact with each other in eco-evolutionary feedback? Our current work provided a valuable starting point for answering those new questions. We encourage future empirical and theoretical studies to go beyond considering parental care as a unitary trait and delve deeper into its components, such as different forms and stages across a breeding cycle and throughout life.

## Methods

### Sex roles classification

We surveyed all 1533 bird species in the Birds of the World database^46^ for which sex provides parental care in each of the three forms –– nest building, incubation, and offspring provisioning –– across a reproductive cycle. We selected the three forms because of the affluence of data and the representative of parental care. We took notes of the parental care features for each species from the breeding section of the species’ account and then classified them into four categories for each form of care: (1) ‘Male care’, where only paternal care was present; (2) ‘Female care’, where only maternal care was present; (3) ‘Biparental care’, where both parents provide care, and (4) ‘Cooperation’, where helpers of cooperatively breeding species also participate in caring of offspring (typically offspring provisioning). Since we are interested in the overall patterns across different bird species at an evolutionary scale, we did not consider the within-species variations of sex roles in this study. Therefore, cases where a form of care was provided usually by females alone but males were occasionally observed to participate were classified as ‘Female care’, and vice versa. In rare cases (N=49 species), the parental care information was recorded with uncertain words, such as “reportedly” or “probably” in one or more care forms (e.g., White-throated Bulbul: nest reportedly built by both sexes; … incubation possibly by both sexes, period 13 days; chicks fed by both parents). All statistical models were run by first including and then excluding those uncertain data.

In particular, for precocial species in which the young are relatively mature and mobile from hatching (i.e., the young leaves the nest shortly after hatching), although parents usually do not feed the precocial chicks, they still invest intensive care efforts (e.g., leading chicks to the food) until their offspring’s independence. In this case, we classified the provisioner sex as the sex who cared for chicks before independence.

In some species of cooperative breeding, the sex role categorization in each care form was straightforward (e.g. White Helmet-shrike: Cooperative breeder, all group-members assisting in all aspects of nesting duties. Breeding pair chooses nest-site and does most of the construction work, but is assisted by other group members; incubation by all group-members; chicks are brooded and fed by all of the group). In the others, the contribution of helpers to each care form may not be specified. Given that cooperative breeding with helpers usually implies helpers’ participation in chick provisioning^47^, we classified those species’ offspring provisioning as ‘Cooperation’, and classified the other two care forms according to additional details in the description regarding sex roles. For example, according to the description “Drakensberg Rockjumper: breeds as monogamous pair and co-operative, with helpers. Nest built by both sexes, …; incubation by both sexes; no other information.”, we classified this species’ nest building and incubation as ‘Biparental care’, and offspring provisioning as ‘Cooperation’.

Following the above procedures, we collected 1533 species with ‘full data’ (i.e., information about sex roles in all three care forms; 651 non-passerines and 882 passerine species). We then matched the scientific names used in the data source^46^ with the species names from a phylogenetic information source (BirdTree.org)^84^. We included 1410 species where we have complete data on the phylogenetic information and contributor(s) of parental care in nest building, incubation, and offspring provisioning for further analyses using statistical models.

### Explanatory variables in statistical models

(a) Sexual selection is the PC1 of the mating system, the sexual size dimorphism, and the sexual dichromatism. Specifically, mating systems of passerine species were obtained following Dale et al. (2015)^42^, and we added non-passerine species that were scored following the same principles. In short, the mating system was scored on a seven-point scale, with ‘0’ representing strict social monogamy (e.g. Zebra finch *Taeniopygia guttata*), ‘1’ representing monogamy with infrequent instances of polygyny observed (< 5% of males, e.g. Lazuli bunting *Passerina amoena*) and ‘-1’ representing monogamy with infrequent instances of polyandry observed (< 5% of females, e.g. Northern flicker *Colaptes auratus*), ‘2’ representing mostly social monogamy with regular occurrences of facultative social polygyny (5 to 20% of males, e.g. American redstart *Setophaga ruticilla*) and ‘-2’ representing mostly social monogamy with regular occurrences of facultative social polyandry (5 to 20% of females, e.g. Pale chanting-goshawk *Melierax canorus*), and ‘3’ representing obligate resource defense polygyny (> 20% of males, e.g., Lance-tailed manakin *Chiroxiphia lanceolata*) and ‘-3’ representing obligate resource defense polyandry (> 20% of females, e.g. Comb-crested jacana *Irediparra gallinacea*). A small number of species with polygynandrous mating systems were pooled with the monogamous species (e.g. Dunnock, *prunella modularis*). Sexual size dimorphism was estimated by combining differences between the sexes in adult body mass, tarsus length, and wing length. In practice, sexual size dimorphism was calculated for three traits representing body size (body mass (g), tarsus length (mm) and wing length (mm)) and was calculated as log (male trait value/female trait value)^85^. Sexual dichromatism was obtained following Gonzalez-Voyer et al. (2022)^86^. In short, the mean value of plumage dimorphism is estimated from five body regions (head, back, belly, wings and tail). Plumage was scored using Birds of the World database^46^. The nominate subspecies of each species was scored using plates as the main reference supplemented with images and descriptions. A single observer scored each body part separately using the following scheme: -2, the female was substantially brighter and/or more patterned than the male; -1, the female was brighter and/or more patterned than the male; 0, there was no sex difference in the body region or there was a difference but neither could be considered brighter than the other; 1, the male was brighter and/or more patterned than the female; 2, the male was substantially brighter and/or more patterned than the female. Thus, positive values represent male-biased ornamentation, zero represents unbiased ornamentation, and negative values represent female-biased ornamentation. The average score of five body regions correlated well with three independent datasets of dichromatism: Spearman rank correlations, rs = 0.705, N = 5825 species, p < 0.001^42^, rs = 0.867, N = 905 species, p < 0.001^87^, and rs = 0.542, N = 855 species, p < 0.001^86,88^.

(b) EPP was the proportion of extra-pair offspring (N = 160 species) and (c) EPBr was the proportion of broods with extra-pair offspring (N = 162 species). Data on EPP and EPBr was obtained from the study of Brouwer & Griffith (2019)^89^. (d) Daily nest predation rate (log10 transformed, N = 225 species) was obtained from Matysioková & Remeš (2018)^75^ and Unzeta et al. (2020)^90^. (e) Clutch size (log10 transformed, N = 1270 species) and (g) length of the nestling developmental period (in days, log10 transformed, N = 1041 species) were collated from Cooney et al. (2020)^91^. (h) Research effort (N = 1376 species), quantified as the number of independent entries per species in the Zoological Record database^92^, was incorporated to account for data quality.

### Statistical analyses

All analyses were carried out within R statistical environment^93^. Firstly, we summarized the variation of sex roles across nest building, nest incubation and offspring provisioning. Next, we tested the three hypotheses (i.e., the ‘consistent expertise model’, the ‘complementary negotiation model’, and the ‘distinct pattern model’) regarding the patterns of sex roles across distinct parental care forms. To do this, we estimated both direct phenotypical Pearson correlations and phylogenetic correlations between sex roles of these three reproductive stages. In particular, we implemented a multivariate phylogenetic model from the package ‘MCMCglmm’^94^ to investigate phylogenetic correlations between sex roles. Following the ‘consistent expertise model’, for each species, sex roles should be similar and consistent across three forms of care and thus we expect positive correlations between them. While according to the ‘complementary negotiation model’, sex roles of two consecutive stages should be the opposite. Therefore, we expected a negative correlation of sex roles between the stage of nest building and incubation, followed by a negative correlation between the stage of nest incubation and offspring provisioning. Third, following the ‘distinct pattern model’, instead of either consistent or complementary sex roles across different parental care forms, we expected that the sex roles across three parental care forms to be rather random and independent (i.e., the expected correlations of sex roles between stages would be neither positive nor negative in this case). Finally, we tried to uncover possible driving forces of the variation of sex differences in parental care across different care forms. This was done by using linear mixed-effect models from the package ‘lme4’^95^ and phylogenetic regression models from the package ‘phylolm’^96,97^.

### Two distinct ways to recode the response variables

Considering the differences in premises regarding different hypotheses and the number of species available for relevant explanatory variables, we coded the response variables (i.e. the contributor(s) of parental care in each form) in two different ways, depending on the corresponding explanatory variables in a series of models. The first way of recoding the contributor(s) of parental care focuses on which sex provides the care. We recoded ‘Female care’, ‘Biparental care’ and ‘Male care’ as ‘-1’, ‘0’, and ‘+1’, respectively. Species in the ‘Cooperation’ category were also coded as ‘0’, because breeders and helpers of both sexes contributed to care. Using this way of recoding, we built five models to test hypotheses regarding the variation/correlations of sex roles across stages of a breeding cycle and investigate whether sexual selection, extra-pair paternity, and nest predation were the main driving factors determining which sex provides care in each form. The second way of recoding the contributor(s) of parental care focuses on the number of individuals that provide care to a brood in each of the three forms. In this way, we recoded ‘Female care’ and ‘Male care’ as ‘1’, ‘Biparental care’ as ‘2’, and ‘Cooperation’ as ‘3’, since there was one carer (either the male or the female) in the first category, two carers (both the male and female parent) in the second category, and at least three carers (both the male and female breeder and at least one helper) in the third category. The second way of recoding allowed us to build an additional model to test the association between the offspring’s life history traits (reflecting the offspring’s reproductive value and brood needs) and the number of carers in each care form.

### Multivariate phylogenetic model estimating phylogenetic correlations

We implemented the multivariate phylogenetic model using the package ‘MCMCglmm’^94^. In this model, sex roles of three parental care forms (nest building, nest incubation and offspring provisioning) were added as three response variables. We first randomly selected a tree of species from the BirdTree^84^ to include in the multivariate model. This tree of 1050 species was then inversed into a phylogenetic covariance matrix (n = 110, 2500 elements) and added as a random effect (similar to including a pedigree matrix as a random effect in an ‘Animal Model’ in quantitative genetics). For the fixed effects, we included ‘sexual selection’, and ‘research effort’. The three-trait MCMCglmm model was set with a proper prior with all variances set to 0.02, covariances set to zero, and a degree of belief parameter set to ? = (size of the matrix +1) = 4. After a burn-in of 10,000 iterations, we ran 260,000 iterations from which a total of 250 samples were drawn (every 1,000 iterations). The high thinning interval was required to eliminate temporal autocorrelations between samples. The large phylogenetic covariance matrix and running iterations indicate this model is challenging in terms of the number of parameters to be estimated. By using these multivariate phylogenetic models, we aimed to estimate the ‘heritability’ (i.e., the ‘phylogenetic signal’ in this case) of each response variable and the ‘genetic correlations’ (i.e., the ‘phylogenetic correlations’ in this case) between the three response variables for further testing the three hypotheses (i.e., the ‘consistent expertise’, ‘complementary negotiation’, and ‘distinct pattern’ models) regarding the relationship of sex roles from the stage of nest building to the stage of offspring provisioning.

### Linear mixed-effect models to uncover possible driving forces

To reveal shared intrinsic drivers regarding the pattern of sex roles across different care stages, we implemented five linear mixed-effects models in addition. Detailed information about the five models is listed below.

**Model 1**: The model was built to quantify the association between sex roles in each of the three forms of parental care and sexual selection. In this model, the sex roles of parental care (using the first way of recoding) were added as the response variable. We included ‘form of care’ (three levels: nest building, incubation, and offspring provisioning), ‘sexual selection’, and ‘research effort’ as fixed effects. The ‘family’ and ‘genus’ of the species were included as a random effect to account for phylogenetic uncertainty. To control phylogenetic uncertainty, we additionally used a phylogenetically controlled regression method as implemented in the function ‘phylolm’ from the R package phylolm^96,97^. In this model, we included the phylogenetic tree of species as a random effect, and ran the model using 100 different phylogenetic trees^84^. Results are therefore based on mean estimates for predictor slopes and model-averaged standard errors. However, because our dataset included multiple traits for each species (i.e. sex roles in nest building, nest incubation and offspring provisioning) and the three different traits were considered as ‘repeated values’ in the model, directly adding a phylogenetic tree into the model as a random effect might be unattainable. To solve this issue, we randomly selected one representative observation (out of three) for each species and ran the model with the culled data, which contained only one observation per species. Note that in this way, the full control for phylogeny was achieved at the cost of losing two-thirds of the observations.

**Model 2 and Model 3**: The two models were built to assess the association between sex roles in each of the three forms of parental care and the degrees of uncertainty in paternity. Sex roles of parental care (using the first way of recoding) were added as the response variable. We included ‘form of care’, ‘body size’, ‘research effort’, and either ‘EPP’ (in Model 2) or ‘EPBr’ (in Model 3) as fixed effects. Regarding random effect(s), due to the limited number of species with data on extra-pair paternity (see the section of ‘explanatory variables’ above), the model simultaneously contained two random effects (‘family’ and ‘genus’) that caused a singular fit issue. This indicated model overfitting, meaning that the random effects structure was too complex to be parameterized by the limited data. Therefore, we only included one random effect (either ‘family’ or ‘genus’) that explained more variation of the response variable. Additionally, we built two phylogenetically controlled regression models using the same fixed effects but treated the phylogenetic tree as a random effect using the phylolm package as explained in Model 1.

**Model 4**: The model was built to test the association between sex roles in each of the three forms of parental care and daily nest predation rates. Sex roles of parental care (using the first way of recoding) were added as the response variable. We included ‘daily nest predation’, ‘form of care’, ‘body size’, and ‘research effort’ as fixed effects, and ‘family’ and ‘genus’ of the species as random effects. Additionally, we built a complementary phylogenetically controlled regression model with the same fixed effects while using the phylogenetic tree as a random effect.

**Model 5**: The model was built to test the association between the number of carers in different care forms and the offspring’s life history traits. In this model, sex roles of parental care (using the second way of recoding) were added as the response variable. We included ‘form of care’, ‘length of nestling developmental period’, ‘clutch size’, and ‘research effort’ as fixed effects. Like in the other models, the ‘family’ and ‘genus’ of the species were treated as random effects to account for phylogenetic uncertainty. Additionally, like in Models 1 to 4, we built a complementary phylogenetically controlled regression model using the phylogenetic tree as a random effect.

## Acknowledgements

We thank Yuansi He and Xinyi Jiang for drawing nice silhouette figures.

