## Supplemental Tables for "Who takes care of the kids at when? Sex differences in avian parental care"

Supplementary Table S1. Output summary of multivariate phylogenetic model estimating phylogenetic correlations. Here, the species tree was inversed into a phylogenetic covariance matrix and added as a random effect. The ‘phylogenetic signal’ of each response variable (grey background) and ‘phylogenetic correlations’ between these three response variables were shown.

| Effect | | Nest building | Incubation | Offspring provisioning |
|---|---|---|---|---|
| genetic | Nest building | 0.800 ± 0.021 | 0.509 ± 0.051 | 0.331 ± 0.055 |
| | Nest incubation | | 0.902 ± 0.016 | 0.419 ± 0.050 |
| | Offspring provisioning | | | 0.769 ± 0.026 |
| Residual | Nest building | 0.200 ± 0.021 | | |
| | Nest incubation | | 0.098 ± 0.016 | |
| | Offspring provisioning | | | 0.231 ± 0.026 |

Table 2. Summary of statistics of five phylogenetically controlled regression models (Model 1 to Model 5). For the random effect (i.e., the phylogenetic tree), the estimated λ is shown. The estimate with its standard error (SE), $t$-value, and corresponding $p$ value is shown for each fixed effect. Note that, for each model, we ran the model using 100 different phylogenetic trees from Jetz et al. (2012). Results are therefore based on mean estimates for predictor slopes and model-averaged standard errors.

| Model 1 | | Estimate (β ± SE) | t | P |
|---|---|---|---|---|
| | Random effects: | | | |
| | Phylogeny (λ) | 0.59 (0.47 – 0.67) | | |
| | Fixed effects: | | | |
| | Intercept | -0.031 ± 0.157 | -0.2 | - |
| | Nest incubation | -0.178 ± 0.029 | -6.2 | <0.001 |
| | Offspring provisioning | 0.274 ± 0.029 | 9.6 | <0.001 |
| | Sexual selection | -0.103 ± 0.013 | -7.9 | <0.001 |
| | Research effort | 0.00002 ± 0.00004 | 0.5 | 0.59 |
| Model 2 | Random effects: | | | |
| | Phylogeny (λ) | 0.91 (0.77 – 1) | | |
| | Fixed effects: | | | |
| | Intercept | -0.058 ± 0.279 | -0.2 | - |
| | Nest incubation | -0.112 ± 0.064 | -1.8 | 0.08 |
| | Offspring provisioning | 0.396 ± 0.075 | 5.3 | <0.001 |
| | EPP | -0.006 ± 0.002 | -2.7 | 0.01 |
| | Research effort | -0.00003 ± 0.00006 | -0.5 | 0.60 |
| Model 3 | Random effects: | | | |
| | Phylogeny (λ) | 0.88 (0.72 – 0.98) | | |
| | Fixed effects: | | | |
| | Intercept | -0.046 ± 0.275 | -0.2 | - |
| | Nest incubation | -0.114 ± 0.066 | -1.7 | 0.09 |
| | Offspring provisioning | 0.433 ± 0.075 | 5.8 | <0.001 |
| | EPBr | -0.002 ± 0.001 | -1.5 | 0.15 |
| | Research effort | -0.00001 ± 0.00006 | -0.2 | 0.82 |
| Model 4 | Random effects: | | | |
| | Phylogeny (λ) | 0.58 (0.28 – 0.76) | | |
| | Fixed effects: | | | |
| | Intercept | -0.565 ± 0.148 | -3.8 | - |

| | | | | |
|---|---|---|---|---|
| | Nest incubation | -0.334 ± 0.055 | -6.0 | <0.001 |
| | Offspring provisioning | 0.484 ± 0.059 | 8.2 | <0.001 |
| | Nest daily predation rate | 0.788 ± 1.262 | 0.6 | 0.53 |
| | Research effort | -0.0001 ± 0.00007 | -1.3 | 0.19 |
| Model 5 | Random effects: | | | |
| | Phylogeny (λ) | 0.68 (0.58 -0.75) | | |
| | Fixed effects: | | | |
| | Intercept | 1.521 ± 0.345 | 4.4 | - |
| | Nest incubation | -0.172 ± 0.033 | -5.3 | <0.001 |
| | Offspring provisioning | 0.326 ± 0.033 | 10.1 | <0.001 |
| | Nestling developmental time | 0.084 ± 0.145 | 0.6 | 0.56 |
| | Clutch size | -0.076 ± 0.057 | -1.3 | 0.18 |
| | Research effort | -0.00004 ± 0.00004 | -0.9 | 0.35 |

Supplementary Table S3. Output summary of five linear mixed-effects models (Model 1 to Model 5) by excluding uncertain species. For random effects, the size of the variance component was shown. The estimate with its SE, $t$ value, and corresponding $p$ value was shown for each fixed effect.

| Model 1 | | Estimate (β ± SE) | t | P |
|---|---|---|---|---|
| | Random effects: | | | |
| | Family (n = 128) | 0.063 | | |
| | Genus (n = 470) | 0.017 | | |
| | Residual | 0.127 | | |
| | Fixed effects: | | | |
| | Intercept | -0.298 ± 0.028 | -10.7 | - |
| | Nest incubation | -0.181 ± 0.016 | -11.5 | <0.001 |
| | Offspring provisioning | 0.264 ± 0.016 | 16.8 | <0.001 |
| | Sexual selection | -0.107 ± 0.009 | -11.3 | <0.001 |
| | Research effort | 0.00005 ± 0.00002 | 2.3 | 0.03 |
| Model 2 | Random effects: | | | |
| | Family (n = 64) | 0.109 | | |
| | Genus (n = 124) | 0.006 | | |
| | Residual | 0.140 | | |
| | Fixed effects: | | | |
| | Intercept | -0.256 ± 0.060 | -4.3 | - |
| | Nest incubation | -0.182 ± 0.042 | -4.3 | <0.001 |
| | Offspring provisioning | 0.346 ± 0.042 | 8.2 | <0.001 |
| | EPP | -0.004 ± 0.002 | -1.8 | 0.07 |
| | Research effort | -0.000001 ± 0.00005 | -0.2 | 0.79 |
| Model 3 | Random effects: | | | |
| | Family (n = 65) | 0.112 | | |
| | Genus (n = 125) | 0.006 | | |
| | Residual | 0.139 | | |
| | Fixed effects: | | | |
| | Intercept | -0.271 ± 0.060 | -4.5 | - |
| | Nest incubation | -0.193 ± 0.042 | -4.6 | <0.001 |
| | Offspring provisioning | 0.329 ± 0.042 | 7.9 | <0.001 |
| | EPBr | -0.0007 ± 0.001 | -0.6 | 0.53 |
| | Research effort | 0.000 ± 0.000 | -0.1 | 0.89 |
| Model 4 | Random effects: | | | |
| | Family (n = 55) | 0.033 | | |
| | Genus (n = 138) | 0.011 | | |
| | Residual | 0.121 | | |

| | | | | |
|---|---|---|---|---|
| | Fixed effects: | | | |
| | Intercept | -0.500 ± 0.044 | -11.3 | - |
| | Nest incubation | -0.241 ± 0.033 | -7.3 | <0.001 |
| | Offspring provisioning | 0.527 ± 0.033 | 16.0 | <0.001 |
| | Nest daily predation rate | -0.149 ± 0.888 | -0.2 | 0.86 |
| | Research effort | 0.000 ± 0.000 | -0.02 | 0.99 |
| Model 5 | Random effects: | | | |
| | Family (n = 138) | 0.097 | | |
| | Genus (n = 483) | 0.042 | | |
| | Residual | 0.140 | | |
| | Fixed effects: | | | |
| | Intercept | 1.546 ± 0.166 | 9.3 | - |
| | Nest incubation | -0.126 ± 0.017 | -7.3 | <0.001 |
| | Offspring provisioning | 0.341 ± 0.017 | 19.7 | <0.001 |
| | Nestling developmental time | 0.137 ± 0.096 | 1.4 | 0.15 |
| | Clutch size | -0.055 ± 0.038 | -1.4 | 0.15 |
| | Research effort | -0.00003 ± 0.00003 | -1.2 | 0.22 |
